# Cytoplasmic divisions without nuclei

**DOI:** 10.1101/2022.06.15.496343

**Authors:** Anand Bakshi, Fabio Echegaray Iturra, Andrew Alamban, Miquel Rosas-Salvans, Sophie Dumont, Mustafa G. Aydogan

## Abstract

Cytoplasmic divisions have been commonly considered a sequel to nuclear divisions, even in the absence of DNA replication. Here we found in fruit fly embryos that the cytoplasm can compartmentalize and divide without nuclei. Our targeted screen for potential necessary and sufficient conditions revealed that, although the cytoplasmic compartments are tightly associated with centrosomes, they can form without astral microtubules and divide without centrioles. Although a focal pool of microtubules is *necessary* for maintaining cytoplasmic compartments, this is not sufficient for their initial formation. Actin filaments are similarly an essential component of cytoplasmic compartments; however, their myosin II-based contractility is unexpectedly dispensable for divisions. We show that the myosin II-based contractility is instead involved in regulating the *pace* of these divisions. Importantly, our results revealed that the cytoplasmic divisions without nuclei can occur in a periodic manner autonomously of the Cdk-Cyclin oscillator that normally drives the cell cycle. We demonstrate that such autonomy of cytoplasmic divisions is preserved even in normal development, where it is leveraged to extrude mitotically delayed nuclei from the blastoderm, protecting the synchrony of rapid nuclear divisions against local delays in mitotic entry. We propose that an active coordination between otherwise autonomous cycles of cytoplasmic and nuclear divisions acts as a quality control mechanism for genome integrity and partitioning in development.

## Main text

The cell cycle is a series of events that leads to mitotic division, such as the start of centrosome duplication, genome replication, chromosome condensation and spindle formation, followed by cytokinesis (*1, 2*). Prevailing models of mitotic progression have long promoted the idea that the rising levels of Cyclin-dependent kinase 1 (Cdk1) activity (*3, 4*) and/or affinity for substrates (*5, 6*) triggers cell cycle events. Recent work, however, has clearly demonstrated that some events can happen independently of Cdk1 activity (*7, 8*). For instance, centrioles can duplicate autonomously when the cell cycle is halted, both by perturbations in dividing cells (*9–11*) and naturally in non-dividing cells (*12*). Furthermore, DNA replication can continue without cell divisions (i.e., endoreplication) (*13*), and conversely, cells can continue to divide even when DNA replication is inhibited (*14–16*). Strikingly, such divisions without DNA replication can occur even under wild-type conditions, as observed during zebrafish skin expansion (*17*).

Cytoplasmic divisions, on the other hand, have been conceptualized as a sequel to nuclear divisions (*18*). This is the case even for cells that divide without DNA replication, as the nucleus retains its division despite a decrease in hereditary material (*14–17*). What triggers cytoplasmic divisions is speculated to be the mitotic regulation of cell cycle kinases and spindle formation, as they are thought to provide an upstream temporal and spatial control for furrowing, respectively (*19*). To what extent they are *required* for cytoplasmic divisions, however, remains unclear.

The cell cycles in early *Drosophila* embryos are a valuable model system to investigate this question *in vivo*, as the regular spacing of nuclei help partition the blastoderm cortex into membrane-bound energids (i.e., cytoplasmic compartments; see Fig. S1) that divide synchronously (*20, 21*) (Video S1). Each of these compartments is expected to host individual nuclei and associated organelles, and undergo cycles of cytoplasmic divisions that follow the nuclear divisions every 13-15min (*22, 23*). In embryos expressing His2-RFP (nuclei) and MRLC-GFP (cytoplasmic compartments), we confirmed that this is generally the case (Figs. 1A and S1A). In some cases, however, we found an intriguing mismatch between the density of nuclei and the cytoplasmic compartments associated with them.

**Figure 1.**
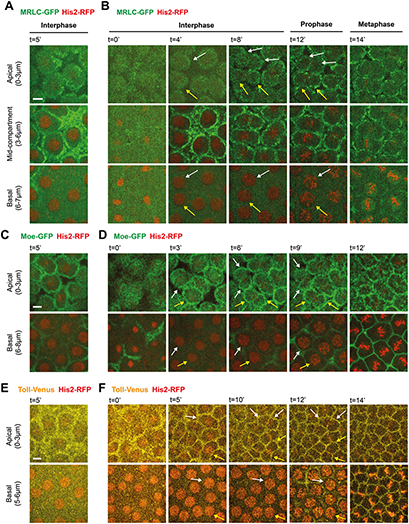
Cytoplasmic compartments can start their divisions during mid-interphase, well prior to mitotic entry, in early fly embryos. Time-lapse micrographs illustrate the progression of interphase and entry into mitosis during cycle 12 or 13 in embryos expressing His2-RFP (marking chromosomes) simultaneously with **(A and B)** MRLC-GFP (myosin), **(C and D)** Moe-GFP (binding actin filaments), or **(E and F)** Toll-Venus (plasma membrane). Top panels (*apical*) visualize the cytoplasmic compartments. Bottom panels (basal) depict the nuclei that reside in each compartment. As expected, cytoplasmic compartments usually host single nuclei **(A, C and E)** and do not begin to divide until after metaphase (see Fig. S1). Surprisingly, however, remaining fraction of cytoplasmic compartments (19.5±6.4%) start dividing in mid/late interphase **(B, D and F)** – see a pair of example compartments (highlighted by *white* and *yellow* arrows) that begin to divide (at the transition from second to third panels in the apical rows). Meanwhile, the nuclei that correspond to these early cytoplasmic divisions (highlighted with matching arrows) remain in interphase without any immediate mitotic entry (B, D and F; see indicated panels in the basal row). Panels are representative examples from embryos expressing indicated transgenes. Percentage data above is represented Mean±SD and calculated from 9 embryos, counting all compartments during Cycle 12 (n≥44 compartments per embryo) from flies that express MRLC-GFP and His2-RFP. Scale bars=5μm.

We noticed that a significant fraction of the cytoplasmic compartments (19.5±6.4%) appeared to start dividing prior to nuclear divisions (5.7±0.9min into interphase 12), or even to mitotic entry, during the blastoderm cell cycles (Fig. 1B). These early divisions occurred in mid-S-phase prior to nuclear envelope breakdown (Fig. S2, A and B). These properties are therefore distinct from the previously identified, constitutively active Rho-A induced myosin-bridges that form during metaphase-to-anaphase transition in these embryos (*24*). To dissect whether these cytokinetic ring-like bridges are mere myosin stripes that bisect the nuclei, or faithful indications of cytoplasmic divisions accompanied by cortex components, we generated flies that express His2-RFP and Moe-GFP (an actin binding molecule). Like MRLC-GFP, Moe-GFP decorated the division rings at the equator of early cytoplasmic divisions (Figs. 1C, 1D and S1B). Similarly, these early divisions were accompanied by plasma membrane ingressions as well, observed in embryos expressing Toll-Venus (a plasma membrane protein) and His2-RFP (Figs. 1E, 1F and S1C). Furthermore, we found that the early cytoplasmic divisions are unlikely to be stochastic events in space and time, as they took place synchronously in mid-to-late interphase and across the whole field of view (Fig. S3, A and B). These results suggested that the early cytoplasmic divisions in fly embryos display some of the quintessential features of normal cell divisions, with the remarkable exception that they can occur synchronously prior to mitotic entry and nuclear division.

Examining the embryos more carefully, we also observed a rare fraction of the cytoplasmic compartments (3.1±2.1%) that, in contrast, were void of nuclei (Fig. 2A— C). Remarkably, these cytoplasmic compartments were fully intact and capable of several rounds of cytoplasmic divisions in the absence of nuclei (Fig. 2A—C; Video S2). These results (Figs. 1 and 2) suggested that the cytoplasmic divisions could occur without nuclei, and hinted at the intriguing possibility that they could be timed autonomously from the mitotic machinery that normally regulate nuclear divisions.

**Figure 2.**
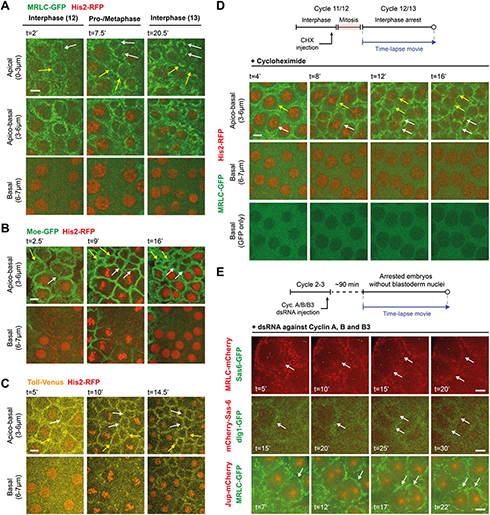
Cytoplasm can divide without nuclei and independently of the principal Cdk-Cyclin oscillator. **(A—C)** Time-lapsed micrographs depict a rare fraction of cytoplasmic compartments (3.1±2.1%) that divide without nuclei in embryos expressing His2-RFP simultaneously with **(A)** MRLC-GFP, **(B)** Moe-GFP, or **(C)** Toll-Venus. Apical or apico-basal panels illustrate the regions where cytoplasmic compartments divide without nuclei in them. Basal panels ensure the lack of nuclei beneath. See a pair of example compartments without nuclei that divide during the cell cycle (highlighted by *white* and *yellow* arrows). Percentage data above is represented Mean±SD and calculated from 9 embryos (the same set as Fig. 1 for comparability), counting all compartments during Cycle 12 (n≥44 compartments per embryo) from flies that express MRLC-GFP and His2-RFP. **(D)** Micrographs illustrate that cytoplasmic compartments can continue to divide (see apico-basal panels), at least for another round, in embryos injected with cycloheximide and arrested in interphase (see basal panels depicting the arrested nuclear cycle and intact nuclear shadows without NEB). Images are a representative set from 4 embryos injected with the drug. **(E)** Cyclin-A-B-B3 triple cocktail dsRNA injection in embryos that express (top panel) MRLC-mCherry and Sas-6-GFP (centrioles); (middle panel) mCherry-Sas-6 and dlg1-GFP (plasma membrane); (bottom panel) MRLC-GFP and Jup-mCherry (microtubules). Illustrated by *white* arrows in each scenario, see the formation and division of compartments without nuclei – judged by the complete lack of nuclear shadows basally, as successfully done before using the nuclear retention of otherwise cytoplasmic proteins (*11*) (see Fig. S4 for validations). Images are representative sets from 9 embryos injected with the dsRNA cocktail. The schemes above panels (D and E) illustrate the injection protocols. Scale bars=5μm.

As mitotic cyclins degrade every cell cycle (*25*), we first sought to determine whether continuous protein synthesis is essential for cytoplasmic divisions, so we injected cycloheximide (a translation inhibitor) into embryos. As expected, the injected embryos were arrested in interphase, evident from the nuclear envelopes that did not break down (Fig. 2D; bottom panel). Despite this dramatic halt to global translation and the cell-cycle progression, the cytoplasmic compartments continued to duplicate for at least another round (Figs. 2D and S3C). Importantly, like the early cytoplasmic divisions that we observed in unperturbed interphases, cytoplasmic divisions in the injected embryos occurred throughout the embryo and did so synchronously (Fig. S3D). This suggested that the synchronous cycles of cytoplasmic divisions are likely reset by post-translational modifications, and raise the possibility that the mechanism regulating their divisions could be distinct from the Cdk-cyclin oscillator that normally governs mitotic progression.

To directly test whether the Cdk-Cyclin oscillator regulates cytoplasmic divisions, we synthesized dsRNA against all mitotic cyclins (A, B and B3) – a method that robustly halts Cdk1 activity and prevents nuclear divisions in fly embryos (*10, 11, 26*). We injected the dsRNA cocktail into embryos that express MRLC-GFP soon after their fertilization (∼cycle 2-4). As centriole duplications can occur independently of nuclear cycle progression (*10, 11*), we simultaneously expressed Sas-6-mCherry to monitor them as a cytological marker, which helped levelling the blastoderm without nuclei. Strikingly, we found that the cytoplasm can compartmentalize without nuclei (Fig. S4) and go through several rounds of divisions periodically (6.4±2.1 min; n=14 compartments from 2 ind. reps.) in arrested embryos (Fig. 2E, top panel; Video S3). Like the cortex of a regular embryo, the cytoplasmic compartments in arrested embryos were ensheathed with plasma membrane (Fig. 2E, middle panel). Likewise, centrioles appeared to have matured into centrosomes, evident from the nucleation of astral microtubules (Fig. 2E, bottom panel), suggesting that these compartments show architectural similarities to their counterparts in embryos that develop normally. Therefore, it is unlikely that cytoplasmic compartments in embryos are just a mere condensation or aggregation of the cytosol around the dividing nuclei. Our findings show that the cytoplasm can compartmentalize and divide without nuclei, and sustain periodic divisions autonomously of the Cdk-cyclin oscillator.

As several intuitive mechanisms were neither necessary nor sufficient for cytoplasmic divisions without nuclei (Supplementary text S1), we performed a screen targeting signalling pathways, cytoplasmic components or enzymes that are important for canonical cell divisions, to test whether their conventional roles are pertinent for the divisions without nuclei in fly embryos.

We began with investigating whether centrioles might be necessary, because centrioles were unanimously present during the early cytoplasmic divisions in mid-interphase (Fig. S5A), the cytoplasmic divisions without nuclei in normal embryos (Fig. S5B), and the autonomous divisions that occur in arrested embryos (Fig. 2E top and bottom panels). It also stands to reason that centrioles might be required, because their duplications in fly embryos, and in other metazoans, can occur independently of nuclear divisions (*9–11*). We generated flies that express MRLC-mCherry and Sas-6-GFP, and laser ablated centrioles. Upon proper ablations judged by the full loss of centriolar signal, we unexpectedly observed that the cytoplasmic compartments remained intact, maintained circularity and continued their division at least for another round despite the lack of centrioles (Fig. 3A; in all four of the successful ablations). This suggested that centrioles are surprisingly not required for these divisions.

**Figure 3.**
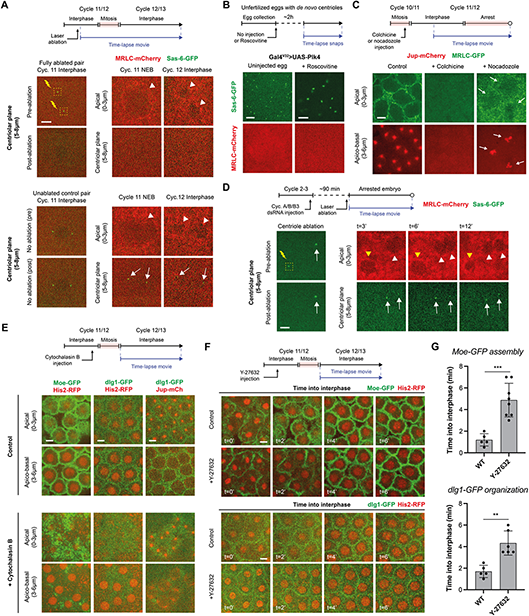
A targeted screen to test potential necessary and sufficient conditions for the formation and division of cytoplasmic compartments in early embryos. **(A)** Micrographs depict compartments (MRLC-mCherry) where the centrioles (Sas-6-GFP) were ablated in early interphase (top set of panels), or were left uninterrupted as a control (bottom set of panels). The success of ablations was judged by the full elimination of the centriolar signal and the persistence in its absence. See the cytoplasmic compartments without centrioles that continue to divide (top set of panels where w*hite* arrowheads indicate these divisions). *White* arrows in bottom set of panels indicate the unablated centrioles as a control. **(B)** Images illustrate *de novo* centriole formation (by Gal^V32^>UAS-Plk4) in unfertilized eggs, either unperturbed (n=5) or injected with a potent Cdk1 inhibitor, Roscovitine (n=9). Neither of these conditions are sufficient to induce the formation of cytoplasmic compartments. See Fig. S6 for time-lapse snapshots of the images displayed here. **(C)** Micrographs depict cytoplasmic compartments, or their absence thereof, in embryos expressing MRLC-GFP and Jup-mCherry under indicated conditions. *White* arrows highlight how cytoplasmic compartments are able to form, though less robustly, also when chromatin mediates focal MT formation. **(D)** Images show laser ablation of centrioles in embryos injected with dsRNA against Cyclin A-B-B3. Time-lapsed *yellow* arrowheads denote a cytoplasmic compartment where the centrioles were ablated. Conversely, time-lapsed *white* arrowheads follow an unperturbed cytoplasmic compartment in these embryos. *White* arrows indicate the unablated control centrioles. **(E)** Micrographs illustrate the state of cytoplasmic compartments in unperturbed embryos (expressing a variety of markers in combination), or in embryos injected with cytochalasin B. Note how the nuclear divisions can continue in the absence of cytoplasmic compartments (bottom set of panels), albeit with a number of nuclear defects (see Fig. S8 and Supplementary text S2). **(F)** Images depict the formation of cytoplasmic compartments at the beginning of interphase (cyc. 12) either in unperturbed embryos (expressing actin-binding or plasma membrane markers simultaneously with a nuclear marker) or in embryos injected with a small-molecule inhibitor of Rho-kinase, Y-27632. Note the delay in compartment formation in Y-27632 injected embryos. **(G)** Bar charts quantify the compartment formation delays depicted in (F). Data are presented as Mean±SD, where each point represents a single embryo. Statistical significance was assessed using a Welch’s t-test (for Gaussian distributed data) or a Mann-Whitney test (***, p<0.001; **, p<0.01). The schemes above panels (A—F) illustrate the injection protocols. Scale bars=5μm.

Although centrioles are dispensable for cytoplasmic divisions, they might be required for the initial formation of cytoplasmic compartments. Previous studies have demonstrated that overexpressing Plk4 can drive *de novo* centriole biogenesis and maturation in fly eggs (*27, 28*). To test this possibility in unfertilized eggs, we first checked whether the cytoplasmic compartments might self-organize by default. Unlike in the homogenized extracts of *Xenopus* eggs (*29*), we found that this is not the case in flies (Fig. S6A). Next, we collected virgin mothers that lay eggs producing thousands of *de novo* centrioles. Importantly, we found that *de novo* centrioles do not trigger the formation of cytoplasmic compartments (Figs. 3B and S6B). As Cdk1 activity is naturally high in eggs, which is refractory for myosin localization (*30*) and might prevent cytoplasmic compartment formation, we injected Roscovitine – a drug that inhibits Cdk1 activity in fly eggs (*31*). Just as the uninjected counterparts, the injected eggs with *de novo* centrioles did not display any cytoplasmic compartments either (Figs. 3B and S6C). These results strongly indicate that neither the centrioles are sufficient to induce cytoplasmic compartments, nor is their duplication/presence necessary for cytoplasmic divisions.

Centrosomes are a major source of astral microtubule (MT) polymerization; however, it is well-appreciated that other MT pools exist (*32*), so we tested whether MT polymerization is required for the formation of cytoplasmic compartments. When we injected colchicine (an inhibitor of global MT polymerization) into embryos expressing MRLC-GFP and Jupiter-mCherry (MTs), the cytoplasmic compartments were heavily impaired (Figs. 3C and S7, A and B). Just as in vertebrate tissues, the redundancy between the astral and chromatin-mediated MT pools is well characterized in fly embryos (*33*), so we next administered low-dose nocodazole to achieve a partial interference on astral MTs (*34, 35*). As expected, while the astral MT capacity was greatly diminished, the chromatin-mediated MT pool appeared to be intact and was promoted even more robustly (Fig. S7C). Despite an early mitotic arrest in response to the injection (leading to transient myosin delocalization), the cytoplasmic compartments formed and remained intact surrounding these MTs (Figs. 3C and S7C). Our results suggest that, a focal pool of MTs, but not centrosomal MTs or a mitotic spindle *per se*, are essential for the formation of cytoplasmic compartments.

Our laser ablation experiments showed that centrioles are not required for cytoplasmic divisions (Fig. 3A). We reasoned that this could be due to a potential “rescue” by the chromatin-mediated MTs under normal conditions. If this were true, ablating the centrioles in cell-cycle arrested embryos – which largely lack nuclei – would seize the autonomous cytoplasmic divisions. Indeed, ablating centrioles in arrested embryos seized cytoplasmic divisions (Fig. 3D; in all five of the successful ablations in arrested embryos). By these findings, we learn that the well-known redundancy in cellular MT organization not only serves to maintain spindle robustness (*33*), but could also help sustain other aspects of cytoplasmic organization, such as the cytoplasmic divisions in embryos as we demonstrate here.

Previous work has suggested that actin filament network is similarly a necessary structural component of cytoplasmic compartments (*22, 36*). To confirm this, we disrupted the actin filament network by a cytochalasin B injection (Video S4 and Fig. 3E; left column). The cytoplasmic compartments were indeed abolished, judged by the absence of an organized plasma membrane (Fig. 3E, middle column). Surprisingly, the nuclear cycles continued relatively normally albeit with several karyotype damages (Video S4, Fig. S8 and Supplementary text S2). Importantly, the cytoskeletal disruption by cytochalasin B was specific to the actin network, as the astral MTs appeared relatively normal (Fig. 3E; right column). This result supports our findings that, although a focal pool of MTs is necessary (Figs. 3C and S7), this *alone* cannot be sufficient for the formation and maintenance of cytoplasmic compartments (Figs. 3B and S6).

Next, we wondered whether actin’s myosin II-based contractility is similarly important. We found by several independent methods that RhoA, an upstream Rho-GTPase effector of myosin, is not required for the formation and divisions of cytoplasmic compartments (Fig. S9, A and B, and Supplementary text S3). Downstream in the same pathway, myosin is regulated via the activating phosphorylations by Rho-associated protein kinase (ROCK). We therefore injected Y-27362, a selective inhibitor of ROCK, preventing myosin II-based contractility in fly embryos (*30*). While Y-27362 significantly impaired cortical myosin recruitment (Fig. S9C), we observed that the cytosolic MRLC-GFP signal appeared to generate subtle halo-like patches around the nuclei (Fig. S9C; see bottom panels at t=4’ and t=6’), hinting that the cytoplasmic compartments may still be intact. Indeed, embryos injected with Y-27362 still displayed intact cytoplasmic compartments decorated with actin filaments and plasma membrane (Fig. 3F). Strikingly, however, we found that the duplication of compartments occurred significantly slower than the wild-type conditions (∼3 min longer than usual, i.e. ∼25% later in the entire interphase duration; Fig. 3, F and G). Together, our results indicate that, although myosin II-based contractility is dispensable for the formation of cytoplasmic compartments and their ability to divide autonomously (Fig. S9, B and D; see Supplementary texts S3 and S4), it regulates the *pace* of cytoplasmic divisions to maintain synchrony with the divisions of nuclei.

Our findings imply that the formation, and timely divisions, of cytoplasmic compartments is essential to sustain proper nuclear spreading, thereby protecting the genome integrity (see Supplementary text S2). This notion is intuitive and strongly supported by the work of others, however it only considers cytoplasmic compartments as “passive” walls that partition nuclei in early embryos. It remains unintuitive why the autonomy of cytoplasmic divisions should be preserved, even in apparently healthy embryos. Unless it confers a physiological advantage, such autonomy would have been rendered redundant in the face of robust cell cycle enzymes through evolution (*7*).

Inspired by this enigma, we examined the real-time relationship between the nuclear and cytoplasmic cycles more carefully. In embryos expressing His2-GFP and MRLC-mCherry, we unexpectedly noticed that the mitotic entry of a very small but reproducible number of nuclei (2.5±1.8 per 100 nuclei; n=9 embryos) was delayed by 1.5-2 min, corresponding to 25-35% of the entire mitosis duration in cycle 12 (Fig. 4A, see the delay of NEB and chromosome condensation in bottom panel; Video S5, *white* arrows labelled “Type 1”). Examining these mitotically delayed nuclei in comparison to their non-delayed, immediate neighbours (Fig. 4A, top panel), we found that the delayed nuclei were – without exception – encapsulated by a cytoplasmic compartment that has already divided in interphase (Fig. 4A, *white* arrows; n=16 mitotically delayed nuclei from 9 embryos). Although not all early cytoplasmic divisions had mitotically delayed nuclei (only ∼15% of the early cytoplasmic divisions presented in Fig. 1), the cytoplasmic compartments bisecting the delayed nuclei furrowed significantly deeper (2.0±0.5μm) than the ones without (0.9±0.4μm; *p*<0.0001, Mann-Whitney test). Moreover, the mitotically delayed nuclei failed to divide (à la mitotic slippage; Fig. 4B, see *white* arrowhead in bottom panel). Therefore, in the next cell cycle, the cytoplasmic compartments that had divided above them appeared to contract and push the undivided nuclei basally, leading to their elimination from the blastoderm (Fig. 4B, bottom panel). Importantly, we found that previously reported conditions (*37–39*) for eliminating nuclei with spindle segregation errors (*nuclear fallout*; here referred to as “Type 2”) are unlikely to fully account for the elimination of mitotically delayed nuclei from the blastoderm (Fig. S10, Video S5 and Supplementary text S5).

**Figure 4.**
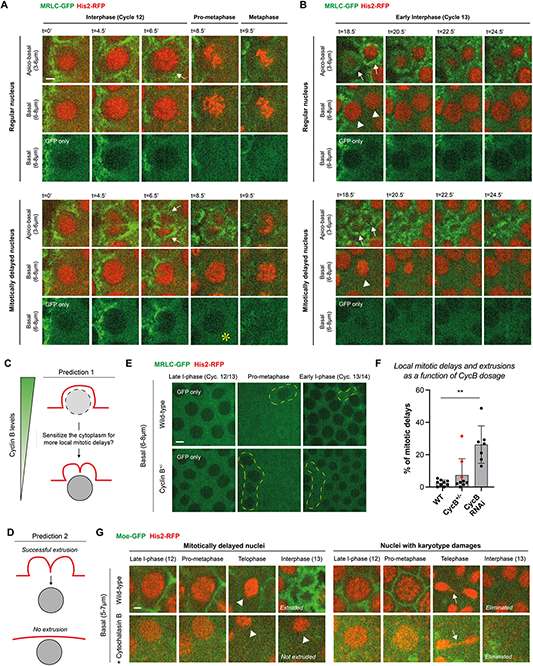
Autonomous cytoplasmic divisions facilitate the extrusion of nuclei that locally delay into mitosis during fly blastoderm formation. **(A and B)** Time-lapse micrographs illustrate the real-time relationship between cytoplasmic divisions (MRLC-GFP) and the nuclear cycles (His2-RFP) both for a nucleus that enters mitosis in time (top set of panels) and one that is delayed into mitosis by 1.5-2min locally (bottom set of panels). Scale bar=2μm. **(A)** Regular “in-time” nuclei are typically in synchrony with their cytoplasmic compartments (*white* arrows; top). In contrast, delayed nuclei are always associated with early cytoplasmic divisions in interphase (*white* arrows; bottom). These delays are evident from several cytological markers, including a visible delay in chromosome condensation and in nuclear envelope breakdown (denoted by a *yellow* asterisk in the basal “GFP-only” panel). **(B)** At the beginning of the next cycle, while regular nuclei have divided and been encapsulated by their compartments (indicated by white arrowheads and arrows, respectively), the delayed nuclei appeared to have slipped mitosis (i.e., skipping nuclear division). Nevertheless, the cytoplasmic compartments that had divided autonomously were intact above the delayed nuclei and appeared to extrude them basally. **(C and D)** Cartoon schematics illustrate two major predictions (referred to as Predictions #1 and #2 in the main text) of our working hypothesis on the mechanism of how mitotic delays are triggered and how the delayed nuclei are eliminated (see Fig. S11 for further details). **(E)** Images show nuclear entry into mitosis in control embryos or in embryos that are heterozygous for Cyclin B (CycB^+/-^), both expressing MRLC-GFP and His2-RFP simultaneously. Mitotically delayed nuclei and their extrusions from the blastoderm are highlighted by *yellow dashed* lines. To emphasize the mitotic delays by virtue of NEBs, only the GFP channel was displayed (see Fig. S12, A and B, for a concomitant depiction of the two channels). Scale bar=5μm **(F)** Bar charts quantify the population percentage of mitotically delayed nuclei and their extrusions in control, CycB^+/-^ or CycB dsRNA-injected embryos expressing MRLC-GFP and His2-RFP (n=9, 9 and 7 embryos, respectively). Data points represent percentage data (Mean±SD) from single embryos where n≥44 nuclei were examined each. 3 *red* data points in the CycB^+/-^ indicate embryos with >5% mitotically delayed nuclei, highlighting that the genetic penetrance of having this response in heterozygote embryos is ∼35%. Statistical significance was assessed using a Kruskal-Wallis test (**, p<0.01). **(G)** Panels illustrate the fate of mitotically delayed nuclei and nuclei with damages either in unperturbed embryos or in embryos injected with cytochalasin B (expressing Moe-GFP and His2-RFP). While in unperturbed embryos both types of nuclei were eliminated from the blastoderm (see Supplementary text S5), the mitotically delayed nuclei (indicated with *white* arrowheads) were no longer extruded in embryos injected with cytochalasin B. The nuclear damages, lagging chromosomes in this case, were denoted by *white* arrows. See main text for sample size and percentage comparisons. Scale bar=2μm.

As such, we propose the existence of a novel blastoderm quality control mechanism in which mitotically delayed nuclei can be *extruded* by the cytoplasmic compartments that divide autonomously and displace them from the blastoderm (Fig. S11). This hypothesis posits two major predictions. Since the elimination of mitotically delayed nuclei is associated with slower NEB and a failure of nuclear division (Fig. 4, A and B), lower levels of Cdk1 activity would be predicted to trigger local mitotic delays more frequently (Fig. 4C; *Prediction 1*). As myosin’s localization to the cortex is normally inhibited by higher Cdk1 activity (*30*), this could also explain why the cytoplasmic compartments that encapsulate mitotically delayed nuclei can divide early and furrow deeper. Second, in the complete absence of cytoplasmic compartments, the delayed nuclei would no longer be extruded (Fig. 4D; *Prediction 2*).

To test whether lowering Cdk1 activity could sensitize the embryos to induce more local mitotic delays (Fig. 4C; Prediction 1), we generated flies that express MRLC-GFP and His2-RFP where the dosage of Cyclin B was halved (CycB^+/-^), and examined their embryos in syncytial cycles (Figs. 4E, S12A and S12B). Excitingly, we found that CycB^+/-^ embryos displayed significantly more local mitotic delays (Fig. S12B and Video S6, encircled with *yellow* dotted lines), accompanied by autonomous cytoplasmic divisions (Fig. S12B, top panels) and followed by clusters of nuclear elimination (Video S6 and Fig. S12B, bottom panels). Since CycB^+/-^ embryos displayed ∼35% genetic penetrance to induce mitotic delays in >5% of all nuclei (Fig. 4F; *red* data points), we sought to test whether a more complete depletion of Cyclin B would elicit higher penetrance. Remarkably, in embryos injected with dsRNA against Cyclin B (10-15min prior to cycle 10), we observed full penetrance (7 out of 7 embryos) in inducing clusters of mitotic delays (Fig. S12C), and do so for ∼25% of all nuclei on average (Fig. 4F). Together, our results strongly indicate that mitotically delayed nuclei and their elimination associate with, and are triggered more frequently, by lower levels of Cdk1 activity.

Finally, we asked whether the cytoplasmic compartments are *required* for the elimination of delayed nuclei (Fig. 4D; Prediction 2), as the cytoplasmic compartments that divide autonomously and furrow deeper appeared to extrude these nuclei. We analyzed nuclear elimination events in embryos injected with cytochalasin B (n=6 embryos), where the cytoplasmic compartments are fully abolished, but the nuclear divisions continue with various karyotype damages (Figs. 3E, S8 and 4G; see Supplementary text S2). Just as the control embryos, the cytochalasin B treated embryos had several nuclei that were delayed into mitosis (Fig. 4G and Video S7, *white* arrows labelled “Type 1”). Remarkably, however, none of the delayed nuclei in these embryos were eliminated (Fig. 4G, n=5 delayed nuclei; Video S7, *white* arrows). Importantly, and like in the control embryos, most of the nuclei with damage (including the occasional micronuclei) continued to be largely eliminated (Fig. 4G, 14 out 19 nuclei with damages were eliminated; Video S7, *yellow* arrows labelled “Type 2”). These results firmly agree with our proposal that the mitotically delayed nuclei are extruded by cytoplasmic compartments that nevertheless divide autonomously, therefore distinct from previously described mechanisms of nuclear elimination induced by chromosome segregation errors (Supplementary text S5).

That the cytoplasm can divide without a nucleus and do so in a periodic manner independently of the Cdk-cyclin oscillator, challenges an important set of assumptions made in the prevailing models of eukaryotic cell divisions. First, not only can cytoplasmic divisions precede spindle formation, they also do not require an intact spindle in the first place. Second, the cytoplasmic divisions can occur twice as fast (∼5-6min) as the nuclear divisions (∼10-15min), suggesting that the Cdk-cyclin oscillator has likely evolved to slow cytoplasmic divisions to align them with cycles of chromosome segregation. This is in contrast to the autonomous cycles of centriole duplications, which occur at a natural period of ∼17min, where nuclear cycles appear to help speed up centriole biogenesis in wild-type conditions (*11*). Together, these findings provide a striking new evidence for the emerging concept of “autonomous clocks” in which the Cdk-cyclin oscillator is postulated to couple otherwise autonomous biological cycles to run at the pace of nuclear cycles (*7, 11, 40, 41*).

Unless it confers a fitness or reproductive advantage, why have flies – species as modern as 250myo – maintained the autonomy of cytoplasmic divisions? Here, we found a previously unreported, minor fraction of nuclei that delay into mitosis and slip division. Despite this local stall in the nuclear cycle, the cytoplasmic compartments divide autonomously and eliminate the undivided nuclei by extrusion. Importantly, lowering Cdk1 activity sensitizes the cytoplasm for even more local mitotic delays and subsequent extrusions. Such nuclear extrusions in the fly blastoderm are reminiscent of a minor fraction of roundworm embryonic cells that also stall progression into mitosis and are subsequently eliminated by an extrusion program (*42*), similarly involving the downregulation of cell-cycle molecules. In another bilaterian, the zebrafish, oncogene-transformed embryonic cells also stall in prophase and appear to be extruded through an unusual division (*43*), where a premature cytokinetic ring separates both the basal and apical parts of the cell from the epithelium, and do this even before the cell can assemble a mitotic spindle. Although the upstream signals that trigger these extrusions may be different, we propose that the mechanisms might converge on a common pathway that preserves genome integrity and partitioning during development.

## Acknowledgements

We thank all members of the Aydogan Laboratory for insightful discussions, as well as Updip Kahlon for her critical read of the manuscript. We are grateful to the laboratories of Daniel Kiehart (Duke) and Patrick O’Farrell (UCSF) for their gifts of Moe-GFP and His2-RFP flies. We thank our neighbours, the members of O’Farrell Laboratory for their remarkable resourcefulness, both technically and conceptually, along the way. The research was funded by NIH R35GM136420 (S.D.), NSF 1548297 Center for Cellular Construction (S.D.), Chan Zuckerberg Biohub (S.D.), Sandler Foundation Investigator Award (7029760; M.G.A.), and UCSF PBBR New Frontiers Research Award (2017078; M.G.A.).

## Author contributions

This study was conceptualised by A.B., F.E.I. and M.G.A. Investigation was done by A.B., F.E.I., A.A., M.R-S. and M.G.A. Data were analysed by A.B., F.E.I., A.A. and M.G.A. Methodology was developed by A.B., F.E.I., M.R-S., S.D. and M.G.A. Project was administrated by M.G.A. Resources were shared/made by A.B., F.E.I., A.A., M.R-S., S.D. and M.G.A. Software work was carried out by A.B., F.E.I., A.A. and M.G.A. Overall supervision was done by M.G.A. Validation experiments/analyses were carried out by A.B. and M.G.A. The main version of this manuscript was drafted by A.B. and M.G.A with significant input from all authors. Finally, A.B., F.E.I., A.A., M.R-S., S.D. and M.G.A. reviewed and edited the manuscript.

## Data and material availability

All microscopy data and experimental materials are immediately available upon request.

## Supplementary Materials

### Materials and methods

#### Fly husbandry, stocks and embryo collections

Flies for experiments were kept at 25°C in *Drosophila* culture medium (7.5% molasses, 1.01% agar, 1.4% agar, 5.6% cornmeal, .75% tegosept, .23% propionic acid, .04% phosphoric acid) in vials or bottles. Stocks were kept in 8x 2.5cm plastic vials. The specific fly alleles used in this study are listed in Table S1. The fly stocks generated and tested in this study are listed in Table S2.

Embryos were collected on juice plates containing 25% cranberry-raspberry juice (2% sucrose and 1.8% agar, with the addition of fresh yeast droplets), and were incubated at 25°C.

#### Microinjection experiments in embryos and unfertilized eggs

Embryos were collected after a 20min incubation with the juice plate, then were aged at 25°C for 50 minutes (so as to begin the injections starting from earlier nuclear cycles, such as cyc. 9-11). After this incubation period, embryos were dechorionated using clear double-sided tape, and mounted on a strip of glue onto a 35mm MatTek glass-bottom petri dish with a 14mm microwell. After desiccation for 6min at 25°C, the embryos were then covered in *Voltalef* oil (Grade H10S). All drug, purified enzyme and dsRNA injections were performed using a borosilicate glass tube, 1.2mm outer diameter, 0.9 mm inner diameter, pulled on a model P-87 micropipette puller. The heat, pull, velocity, and time values were 670, 60, 80, 190 respectively. Uninjected, control embryos were subjected to same treatments except the injection.

Similar to that described by Aydogan et al. 2020, the “early” injection of Cyclin A-B-B3 dsRNA cocktail (but not for the “late” injection of Cyclin B dsRNA only) was performed on embryos that were collected after a 20min incubation time – a method that successfully arrests the nuclear divisions of fly embryos in the earliest cycles it could. For the triple cyclin dsRNA cocktail experiments, the embryos were injected with a final needle concentration of 0.67 mg/ml. For the cyclin B dsRNA experiments, the embryos were injected with a needle concentration of 2 mg/ml. During the injections, all surfaces (including the pipettes) were cleaned with RNase*Zap*^TM^ RNase decontamination solution (ThermoFisher Scientific) and 70% ethanol. Following the injections, the arrested embryos were left to age for 1-1.5h before imaging.

For unfertilized egg experiments, the specimen was collected during a 2h incubation with juice plates at 25°C, then were imaged immediately. For unfertilized eggs where *de novo* centrioles were produced, the specimen was collected during a 2h incubation with juice plates, then were aged at 25°C for another 1-2h before imaging, or before injecting first and imaging afterwards.

#### Drug inventory, purified enzymes and the synthesis of double-stranded RNA

Following drugs were dissolved in nuclease-free water, kept at −20°C and injected at the needle concentration of: Cyclohexamide (50ug/mL; Sigma-Aldrich), Colchicine (100ug/mL; Sigma-Aldrich), Rhosin (19.6mg/mL; R&D Systems) and Y-27632 (76mM; Adipogen). Following drugs were dissolved in DMSO (ranging from 50-100% in stock solution), kept at −20°C, and injected at the needle concentration of: Cytochalasin B (11.7mg/mL; ACROS Organics), Roscovitine (10mM; SelleckChem) and Nocodazole (10mg/mL; Fisher Scientific). Purified Exoenzyme C3 (Cytoskeleton Inc) was prepared with nuclease free water, kept at −80°C and injected at a needle concentration of 100ug/uL.

dsRNAs were synthesized by RNA Greentech LLC (Texas, USA) and stored at −80°C. Coding sequences used to generate the dsRNAs are listed in Table S3. To cross-confirm the validity of the correct RNA product in house, the dsRNAs were ran on electrophoresis (1.5% agarose gel) using 2xRNA loading buffer (ThermoFisher Scientific). For electrophoresis, a 1:1 mixture of loading buffer and dsRNA was heated to 65°C for 5 min and then was transferred to ice to decompose any secondary structure on the dsRNA.

#### Microscopy and image analysis

Live embryos were imaged with a spinning disk confocal system (Perkin Elmer Ultraview) at room temperature, using an Olympus IX70 microscope with a planApo 60X 1.40 oil immersion objective and running the Volocity software. 488 and 561 nm lasers were used to visualize GFP/Venus and RFP/mCherry in multiple fly lines. Using 0.5 um intervals, 20 slices were obtained every 30 seconds. The GFP channel was collected using a 200-350ms exposure, 14.5% laser power, and sensitivity of 140, whereas the RFP/mCherry channel was collected using 300ms exposure, 40.5% laser power, and 127-134 sensitivity. All videos were captured with emission discrimination with corresponding filters.

Images were analyzed in ImageJ-Fiji. Following quantifications were performed in embryos that express MRLC-GFP and His2-RFP, counting the respective incidents manually in embryos: the fraction of cytoplasmic compartments that divided in interphase or that divided without nuclei; the fraction of nuclei that delayed into mitosis; the fraction of nuclei that had karyotype damages. Mitotic delays were judged by the delay of nuclear envelope breakdown (in MRLC-GFP channel) and chromatin condensation (in His2-RFP channel). Nuclear damages were judged largely by assessing whether chromatin took up the phenotypes similar to those listed in Figure S8 (see Supplementary text S2). Both the delay-dependent extrusions and nuclear damage-dependent eliminations were judged by alternating z-slices (see Supplementary text S5).

For early cytoplasmic divisions and nuclear extrusion analyses, furrow depth measurements were performed by alternating Z stacks and recording corresponding values in Excel sheets. To determine relative times for the formation of cytoplasmic compartments in control embryos and in embryos injected with Rho Kinase inhibitor Y-27632, the incidents were determined by visual examinations for when >50% of the compartments were formed in the cytoplasm. A similar approach was taken to judge the time of cytoplasmic divisions in embryos injected with Cycloheximide.

The period of early cytoplasmic divisions (into interphase) was measured by taking an average from multiple compartments in cycle 12 (coming from multiple embryos expressing MRLC-GFP and His2-RFP). The period of compartment divisions without nuclei in arrested embryos was measured by taking an average from division-to-division times for multiple compartments (coming from multiple embryos expressing either MRLC-GFP with His2-RFP, or MRLC-GFP with Jup-mCherry). We note that the initial cytoplasmic divisions in arrested embryos occurred slightly slower (∼10min) than the later ones (∼5-6 min), but due to photobleaching in later parts of the videos (which did not necessarily interfere with our time measurements), the images for display were selected from the initial divisions for ease of visualization.

#### Laser ablation experiments

For laser ablation experiments, samples were imaged using an inverted microscope (Eclipse Ti-E; Nikon) with a spinning disk confocal (CSU-X1; Yokogawa Electric Corporation), head dichroic Semrock Di01-T405/488/568/647 for multicolor imaging, equipped with 405 nm (100 mW), 488 nm (120mW), 561 nm (150mW), and 642 nm (100mW) diode lasers, emission filters ET455/50M, ET525/50M, ET630/75M and ET690/50M for multicolor imaging, and an iXon3 camera (Andor Technology) operated by MetaMorph (7.7.8.0; Molecular Devices)(*44*). Embryos were imaged with a 60x 1.40 Ph3 oil objective.

Laser ablation (30-40 pulses of 3 ns at 20 Hz) with 514 nm light was performed using the MicroPoint Laser System (Andor). Embryos were imaged every 15s with a single stack before ablation, and every 15s with multiples stacks after ablation (0.8μm step size, 11 steps, 8μm total range). Ablating the blastodermal centrioles completely was challenging for several reasons. Centrioles can move laterally in the embryo (even as far and fast as >10um/sec), making it difficult to fully ablate them and limiting our ablation success rates. Also, the centrioles are close (∼5-10μm) to the blastoderm cortex, so slight increases of laser power (to maintain robust ablation) can cause dramatic membrane-injury. Embryos with injured membranes were excluded from analysis. In several cases (4 out of 20 trials in cycling embryos; 5 out of 9 trials in arrested embryos), we were successful in abolishing centrioles without damaging the cortex and drifts in their position. Successful centriole ablations were verified by the immediate elimination of the Sas-6-GFP signal at the site of ablation (centrioles were targeted repeatedly until complete elimination of the signal), and by the subsequent absence of Sas-6-GFP signal recovery in the consecutive timepoints. As demonstrated previously (*45*), the Sas-6-GFP cartwheel structure grows linearly during early/mid interphase in fly embryos, so the dynamics of centriole growth serves as a positive control for proper ablation (failing a proper ablation, the fluorescence would recover within <30sec). Therefore, the centriole ablations were done in early interphase. In case of laser ablations in arrested embryos (injected with the dsRNA cocktail), a similar protocol was followed.

#### Quantification and Statistical Analysis

All measurements in the main text, figures and legends are represented Mean±SD. The details for quantification, statistical significance, sample size, definitions of centers, and the measures for dispersion/precision are indicated in the main body and corresponding figure legends. Statistical significance was defined by p<0.05. To determine distribution normality, data were subjected to D’Agostino-Pearson normality test. GraphPad Prism 9 was used for all statistical analyses.

### Supplementary Text

#### Supplementary text S1

##### The biological rationale for a targeted screen on necessary and sufficient conditions for cytoplasmic divisions in early embryos

In regulating cytoplasmic divisions, our findings exclude a sustenance mechanism based on the *continuous* encoding of an mRNA or cyclic expression of a protein, e.g., the canonical mitotic cyclins that drive nuclear cycles (Fig. 2E). Similarly, another intuitive physical parameter, the major axis length of the compartments, was not significantly different between the ones that divided in interphase (6.2±3.1μm) and the ones that did not (5.2±2.8μm) (n=5 comprt. per N=5 embryos in cyc. 12 per condition; *p*=0.1664, Welch’s test). This result suggests that the cytoplasmic divisions cannot simply depend on the growing size of these compartments. Additionally, we found that neither the nucleus nor translation is required to form and duplicate the cytoplasmic compartments (Fig. 2, A—D). As transcription is also largely silenced in early embryos (*46*) – hence less amenable to *ad hoc* genetic perturbations – we performed a targeted screen composed of biophysical manipulations, maternal mutations, pharmacological inhibitors and purified proteins.

#### Supplementary text S2

##### A detailed assessment of karyotype damages in embryos injected with cytochalasin B

Despite the complete collapse of cytoplasmic compartments in embryos injected with cytochalasin B, several core events of the nuclear cycles continued relatively normally and synchronously post axial expansion (Video S4). At every cycle, the nuclear envelopes broke down, and the chromosomes condensed before they aligned for segregation (Video S4). The segregation morphology was occasionally impaired due to lagging chromosomes, resulting in chromatin bridges and micronuclei – pathologies that are often associated with DNA breakages and/or damages (*47*) (Fig. S8). Also consistent with lagging chromosomes, we observed the formation of synkaryons from metaphase onwards (due to both non-sister and sister nuclei fusion; Fig. S8), likely catalyzed by the uneven spreading and local crowding of nuclei as the cycles advanced (Video S4). These results suggest that, although the formation and duplication of cytoplasmic compartments are clearly not required for nuclear divisions, coupling between the two cycles ensures the genome replication fidelity and helps avoiding abnormal ploidies by spreading the nuclei uniformly prior to morphogenesis.

#### Supplementary text S3

##### RhoA is not required for the formation and divisions of cytoplasmic compartments in early fly embryos

Previous work has suggested that constitutively active RhoA (an upstream Rho-GTPase effector of myosin) can induce myosin bridges in metaphase (*24*). Although these ectopic structures are temporally distinct from the actomyosin bridges we observe in early cytoplasmic divisions (Fig. 1), we reasoned that they might be induced by a similar reaction. As such, we adopted two independent approaches to inhibit Rho-GTPase activity. Exoenzyme C3 transferase (an ADP ribosyl transferase that selectively ribosylates and inhibits RhoA/B/C) inhibits myosin II-based contractility and impairs cellularization starting from the end of cycle 13 in fly embryos (*48, 49*). When we injected cycle 14 embryos with purified recombinant Exoenzyme C3, the cellularization was indeed disrupted (Fig. S9A). When injected in syncytial cycles, however, the cytoplasmic compartments were surprisingly intact, and even showed early interphase divisions as well as divisions without nuclei (Fig. S9B). Similarly, injections of Rhosin hydrochloride (an inhibitor of RhoA’s GEF binding domain) fully mimicked these results (Figs. S9A and S9B). Importantly, in both the Exoenzyme C3 and Rhosin experiments, cortical myosin localization was impaired – in an all-or-none way – only during cellularization (Fig. S9A), but not in the syncytial cycles (Fig. S9B). These suggest that RhoA is not required for the formation and divisions of cytoplasmic compartments in early fly embryos.

That RhoA could be dispensable for cytoplasmic divisions in syncytial fly embryos – but not after the start of cellularization in Cycle 14 – is supported by previous work (*50, 51*), suggesting that the zygotic expression of its canonical effector Rho-GEF (Pebble in *Drosophila*) is required for cytokinesis starting from cycle 14. Although there is evidence that another GEF, Rho-GEF2, is expressed in syncytial embryos (*24, 52*), its partner Rho-GTPase remains unknown. Our findings would strongly suggest that RhoA is an unlikely candidate for the upstream activation of cytoplasmic myosin in these embryos.

#### Supplementary text S4

##### Further consequences of delayed cytoplasmic divisions, as well as their retained ability to divide autonomously, when injected with Y-27362

Besides the visible delay of compartment divisions in embryos injected with Y-27362 (Fig. 3, F and G), the consequences of delay were also manifested in the form of multiple nuclei being encompassed by the same compartment at the beginning of interphase (Fig. 3F), and by the occasional formation of synkaryons due to ill-segregated nuclei (Fig. S9D, right panels). Nevertheless, embryos injected with Y-27362 retained the ability to induce the early cytoplasmic divisions in interphase (though expectedly rarer and mostly in longer interphases such as cycle 13) and the cytoplasmic divisions without nuclei (Fig. S9D; left and middle panels).

#### Supplementary text S5

##### The elimination of mitotically delayed nuclei (*Type 1*) appears distinct from the fallout of nuclei with chromosome segregation errors (*Type 2*)

*Nuclear fallout* is the spontaneous elimination of nuclei from the blastoderm in fly embryos, observed as a consequence of DNA damages that associated with spindle segregation errors, such as lagging and/or bridging chromosomes (*37–39*). Indeed, we also observed that nuclei with these errors were present and, expectedly, were largely eliminated from the blastoderm (Fig. S10, 26 eliminations out of 39 cases with chromosome segregation errors; Video S5, *yellow* arrows labelled “Type 2”). Unlike the mitotically delayed nuclei, the nuclei with segregation errors were neither delayed into mitosis nor failed their mitotic divisions (Fig. S10, A and B; none of the 39 cases examined above). Furthermore, the damaged nuclei were not accompanied by early cytoplasmic divisions that furrowed as deep as the ones associated with the mitotically delayed nuclei (Fig. S10, A and B; none of the 39 cases examined above). In contrast, the mitotically delayed nuclei did not show evidence of damages associated with incomplete DNA replication (Fig. 4, A and B, and Video S5). These results suggest that chromosome segregation errors may not be the sole physiological cause of nuclear eliminations, and that the previously reported conditions for eliminating damaged nuclei are unlikely to fully account for the elimination of mitotically delayed nuclei from the blastoderm.

### Supplementary Tables

**Table S1:**
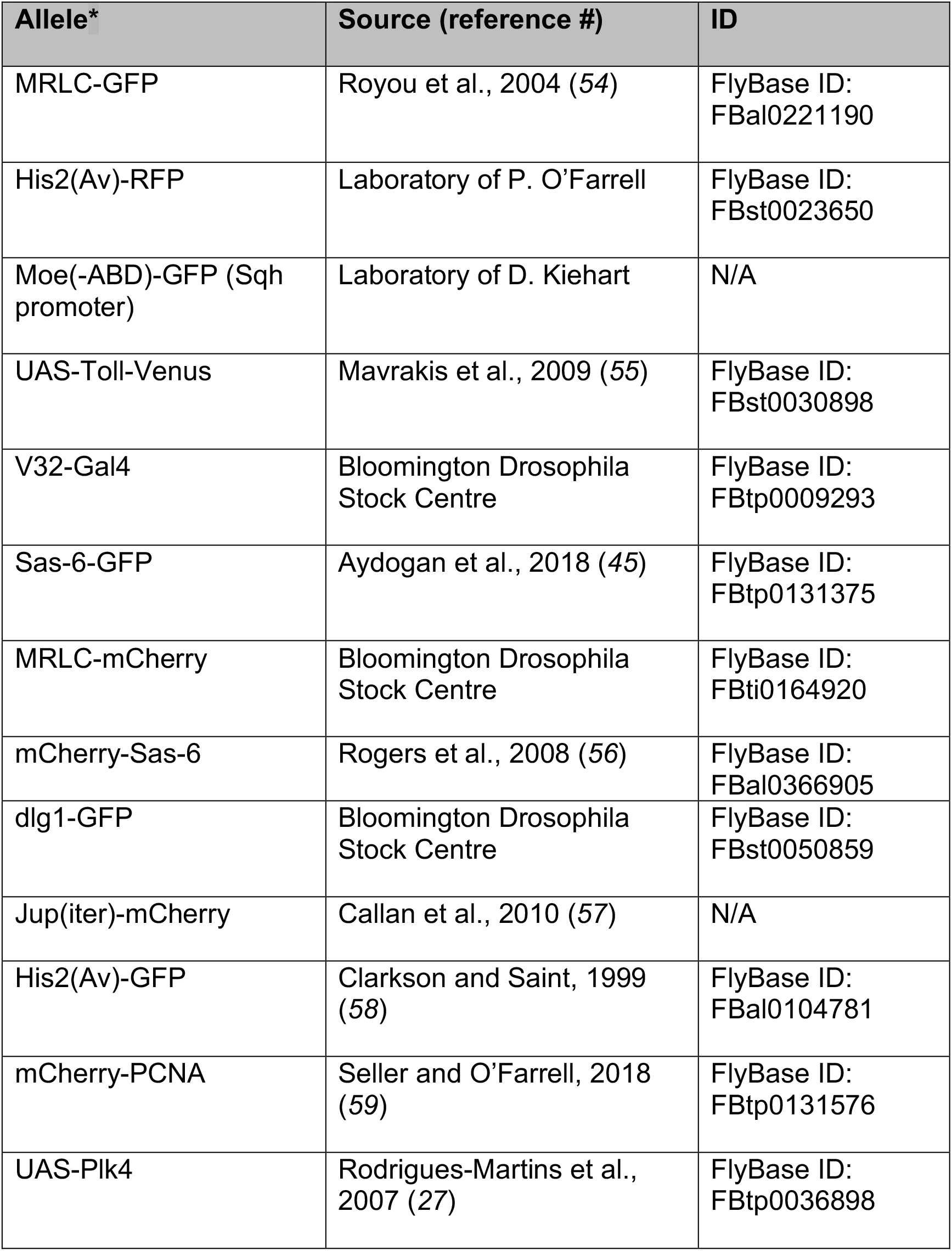
*D. melanogaster* alleles used in this study.

| Allele* | Source (reference #) | ID |
| --- | --- | --- |
| MRLC-GFP | Royou et al., 2004 (54) | FlyBase ID:<br>FBal0221190 |
| His2(Av)-RFP | Laboratory of P. O'Farrell | FlyBase ID:<br>FBst0023650 |
| Moe(-ABD)-GFP (Sqh promoter) | Laboratory of D. Kiehart | N/A |
| UAS-Toll-Venus | Mavrakis et al., 2009 (55) | FlyBase ID:<br>FBst0030898 |
| V32-Gal4 | Bloomington Drosophila Stock Centre | FlyBase ID:<br>FBtp0009293 |
| Sas-6-GFP | Aydogan et al., 2018 (45) | FlyBase ID:<br>FBtp0131375 |
| MRLC-mCherry | Bloomington Drosophila Stock Centre | FlyBase ID:<br>FBti0164920 |
| mCherry-Sas-6 | Rogers et al., 2008 (56) | FlyBase ID:<br>FBal0366905 |
| dlg1-GFP | Bloomington Drosophila Stock Centre | FlyBase ID:<br>FBst0050859 |
| Jup(iter)-mCherry | Callan et al., 2010 (57) | N/A |
| His2(Av)-GFP | Clarkson and Saint, 1999 (58) | FlyBase ID:<br>FBal0104781 |
| mCherry-PCNA | Seller and O'Farrell, 2018 (59) | FlyBase ID:<br>FBtp0131576 |
| UAS-Plk4 | Rodrigues-Martins et al., 2007 (27) | FlyBase ID:<br>FBtp0036898 |

**Table S2:**
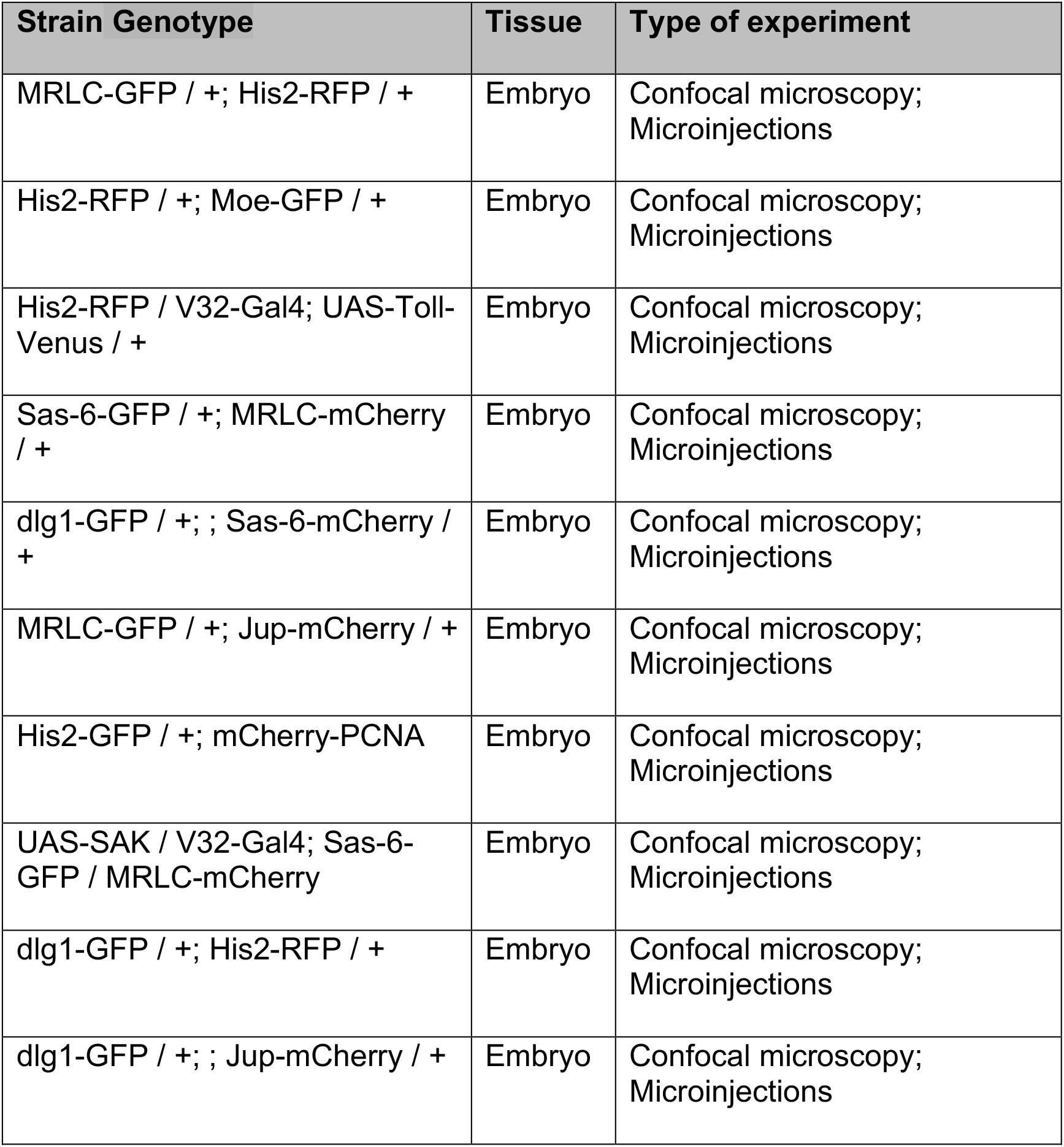
*D. melanogaster* strains generated in this study.

| Strain Genotype | Tissue | Type of experiment |
| --- | --- | --- |
| MRLC-GFP / +; His2-RFP / + | Embryo | Confocal microscopy;<br>Microinjections |
| His2-RFP / +; Moe-GFP / + | Embryo | Confocal microscopy;<br>Microinjections |
| His2-RFP / V32-Gal4; UAS-Toll-Venus / + | Embryo | Confocal microscopy;<br>Microinjections |
| Sas-6-GFP / +; MRLC-mCherry / + | Embryo | Confocal microscopy;<br>Microinjections |
| dlg1-GFP / +; ; Sas-6-mCherry / + | Embryo | Confocal microscopy;<br>Microinjections |
| MRLC-GFP / +; Jup-mCherry / + | Embryo | Confocal microscopy;<br>Microinjections |
| His2-GFP / +; mCherry-PCNA | Embryo | Confocal microscopy;<br>Microinjections |
| UAS-SAK / V32-Gal4; Sas-6-GFP / MRLC-mCherry | Embryo | Confocal microscopy;<br>Microinjections |
| dlg1-GFP / +; His2-RFP / + | Embryo | Confocal microscopy;<br>Microinjections |
| dlg1-GFP / +; ; Jup-mCherry / + | Embryo | Confocal microscopy;<br>Microinjections |

**Table S3:**
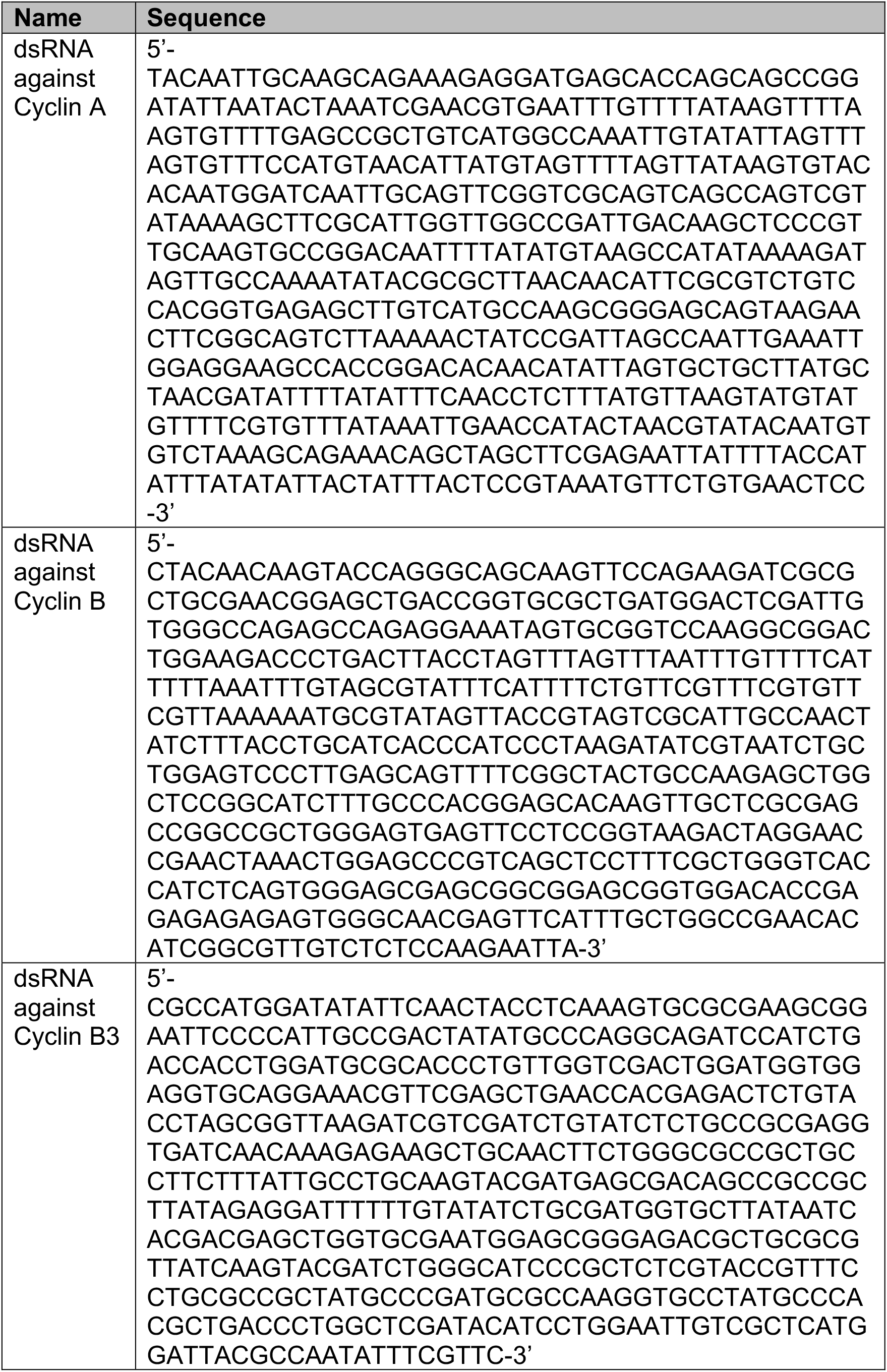
Coding sequences that the dsRNAs were based on in this study.

### Supplementary Figure Legends

**Figure S1.**
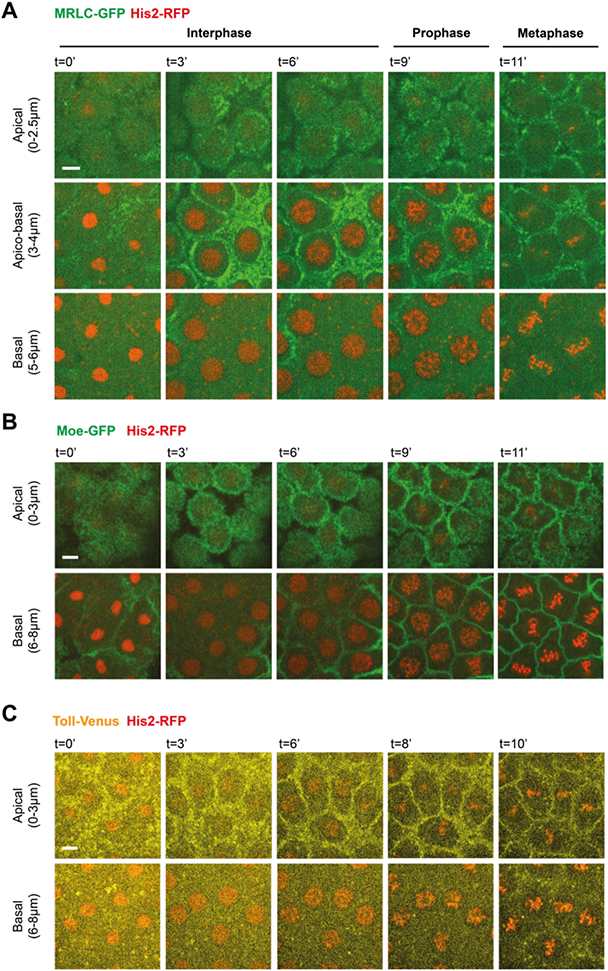
Nuclei in early fly blastoderm are usually hosted within single cytoplasmic compartments, which typically do not divide till the end of mitosis. Micrographs illustrate the progression of interphase and entry into mitosis during cycle 12 or 13 in embryos expressing His2-RFP (marking chromosomes) simultaneously with **(A)** MRLC-GFP (myosin), **(B)** Moe-GFP (binding actin filaments), or **(C)** Toll-Venus (plasma membrane). The panels represent the full time-lapse series of the exact images depicted in Fig. 1A, C and E. Top panels (*apical*) visualize the cytoplasmic compartments (marked by the actomyosin walls and plasma membrane). Bottom panels (basal) depict the nuclei that reside in each compartment. Scale bars=5μm.

**Figure S2.**
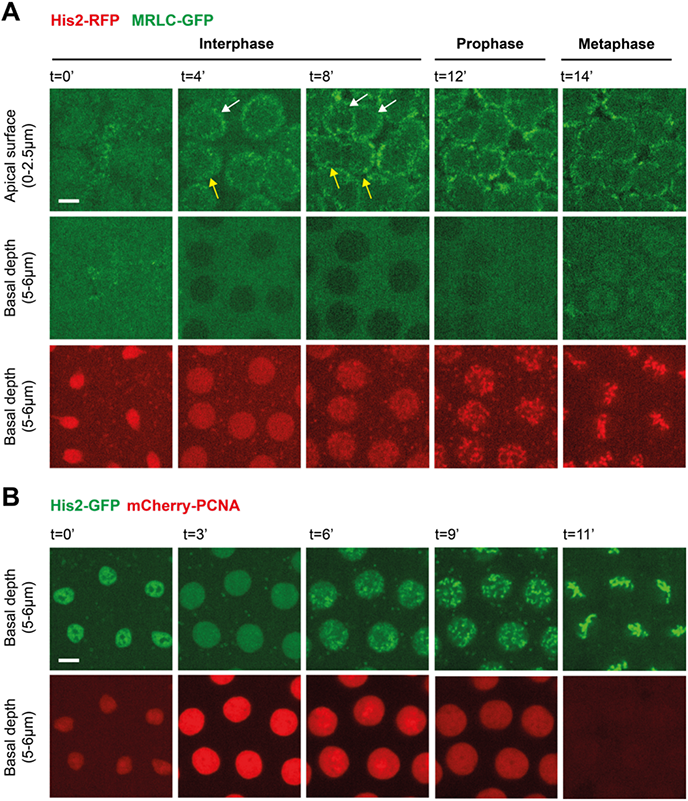
Cytoplasmic divisions can occur well prior to the end of interphase, nuclear envelope breakdown and the start of chromosome condensation. **(A)** Images show the early division of several cytoplasmic compartments in interphase (indicated by *white* and *yellow* arrows), well prior to NEB and chromosome condensation (see mid-panels) in embryos expressing MRLC-GFP and His2-RFP. NEB is judged by the back-illumination of MRLC-GFP. **(B)** Time-lapse micrographs of nuclei correlate the state of His2-GFP and mCherry-PCNA (a common S-phase marker) in order for comparison to the mitotic events that occur in (A). The nuclear retention of mCherry-PCNA indicates that the nuclear envelope is intact. Note that the chromosome condensation in the prophase panel (t=9’) correlates with the start of mCherry-PCNA siphoning, which is fully gone by t=11’, marking the end of interphase and the beginning of mitosis. Scale bars=5μm.

**Figure S3.**
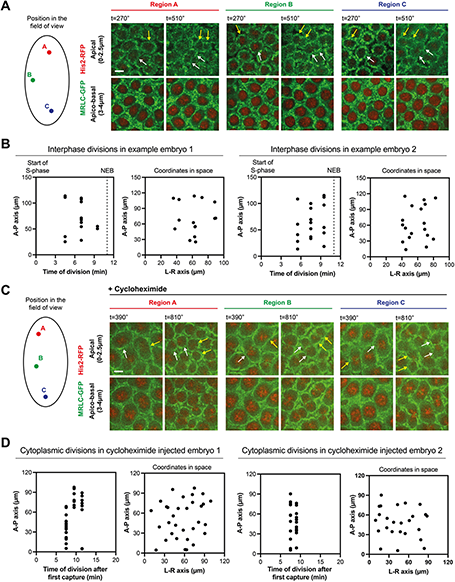
Interphase cytoplasmic divisions occur synchronously in space and time, both in control embryos and in embryos arrested by the injection of cycloheximide. **(A and C)** Micrographs from different regions in the same field of view (denoted with *red*, *green* and *blue* fonts) show cytoplasmic compartments in early interphase (left panels), and their nearly simultaneous start of divisions in space and time (right panels). Representative panels in (A) and (C) are from unperturbed embryos (n=9) and arrested embryos that were injected with cycloheximide (n=5), respectively. Note in cycloheximide injected embryos how the cytoplasmic divisions are more pronounced and furrow even more basally (in comparison to unperturbed embryos). This is presumably because the interphase arrest provides more time for furrowing deeper, as mitoses are normally refractory for this process (*30*). Scale bars=5μm. **(B and D)** In a pair of example single embryos, scatter plots in (B) and (D) quantify the anterior-posterior (A-P) position of interphase cytoplasmic divisions as a function of time during the cell cycle, respectively for (A) and (C). t=0’ marks the start of interphase in (A), and of interphase arrest capture in (C). NEB marks the beginning of mitosis. Similarly, the plots on the right quantify the position of interphase cytoplasmic divisions across A-P and left-right (L-R) axes of embryos, respectively for (A) and (C). Note how interphase cytoplasmic divisions do not cluster into local regions, indicating that a more global program is likely to synchronize these events.

**Figure S4.**
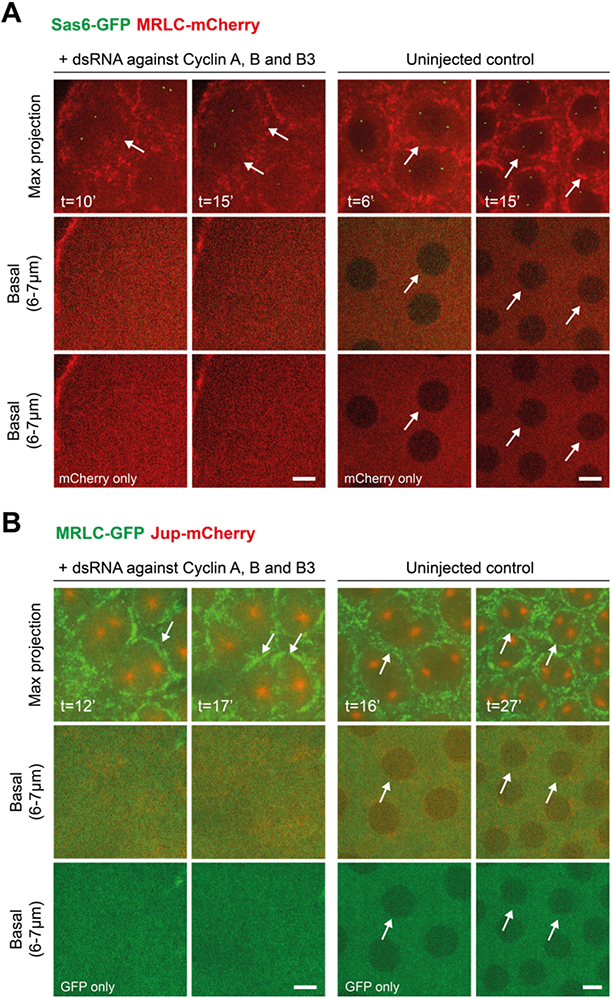
Embryos injected with Cyclins A-B-B3 dsRNA soon after fertilization are largely devoid of nuclei in their blastoderm. Cyclin-A-B-B3 triple cocktail dsRNA injection in embryos that express **(A)** MRLC-mCherry and Sas-6-GFP, and **(B)** MRLC-GFP and Jup-mCherry. Illustrated by *white* arrows in the max-projection panels, see the formation and division of compartments without nuclei. As performed previously using the nuclear retention of Plk4-mNeonGreen and mCherry-Sas-6 (*11*), the lack of nuclei was controlled by examining the back-illumination of MRLC-GFP or MRLC-mCherry channels in basal z-slices, because nuclei normally appear as dark circle shadows in these channels in cycling embryos – see in the *uninjected control* embryos, where additional *white* arrows in basal panels highlight the nuclei that are associated with the cytoplasmic compartments in the top panel. The max-projection images in the dsRNA micrographs are retrieved directly from Fig. 2E for cross-comparison. Scale bars=5μm.

**Figure S5.**
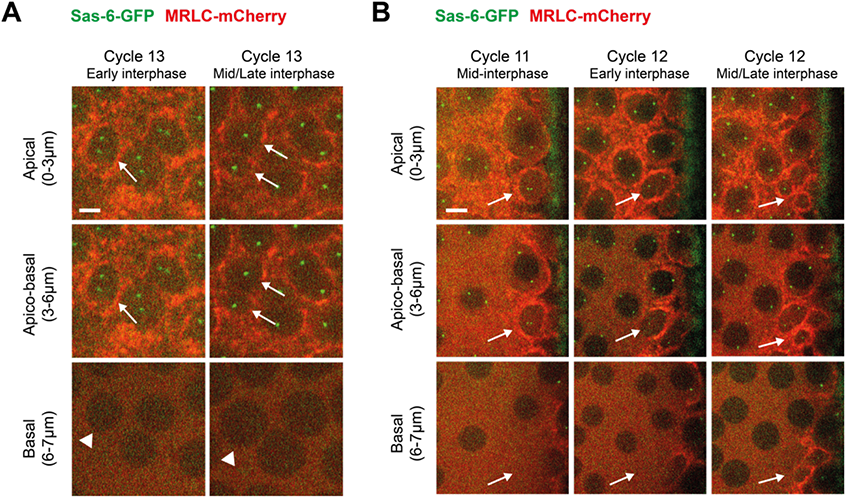
Centrioles are associated with both the cytoplasmic compartments that divide in interphase and the ones that divide without nuclei. **(A)** Images show centrioles (Sas-6-GFP) in a cytoplasmic compartment (MRLC-mCherry) that divides in interphase (indicated by *white* arrows). *White* arrowheads point at the corresponding nucleus that has not entered mitosis. Scale bar=3μm. **(B)** Time-lapse panels show centrioles in cytoplasmic compartments that divide without any resident nucleus. *White* arrows aid following the cytoplasmic divisions apically and the absence of nuclei basally. Scale bar=6μm.

**Figure S6.**
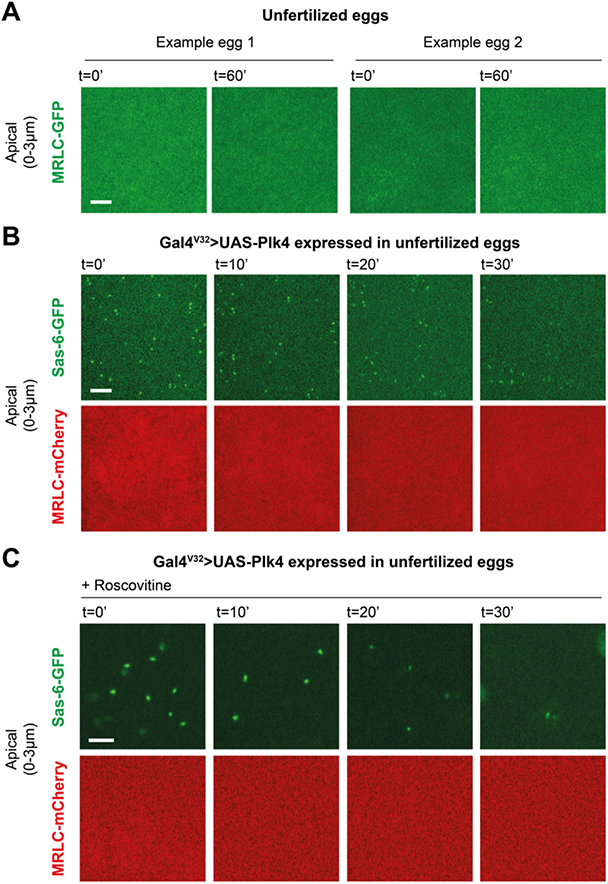
Genetic induction of *de novo* centriole biogenesis is not sufficient to trigger the formation of cytoplasmic compartments in unfertilized eggs. **(A)** Micrographs are obtained from a pair of example unfertilized eggs expressing MRLC-GFP, illustrating that the *Drosophila* egg cytoplasm does not self-organize into compartments by default, even with the passage of time (n=6 embryos). This is in contrast to homogenized *Xenopus* egg extracts that condenses into cell-like compartments *in vitro* (*29*). **(B and C)** Images show that inducing *de novo* centrioles in unfertilized eggs (with Gal4^V32^>UAS-Plk4 expression) is not sufficient to trigger the formation of cytoplasmic compartments, even with the passage of time. This is true both in unperturbed eggs (n=5 embryos) and in eggs injected with Roscovitine (n=8 embryos) to inhibit Cdk1 activity, which is normally refractory for myosin localization (*30*). Scale bars=5μm.

**Figure S7.**
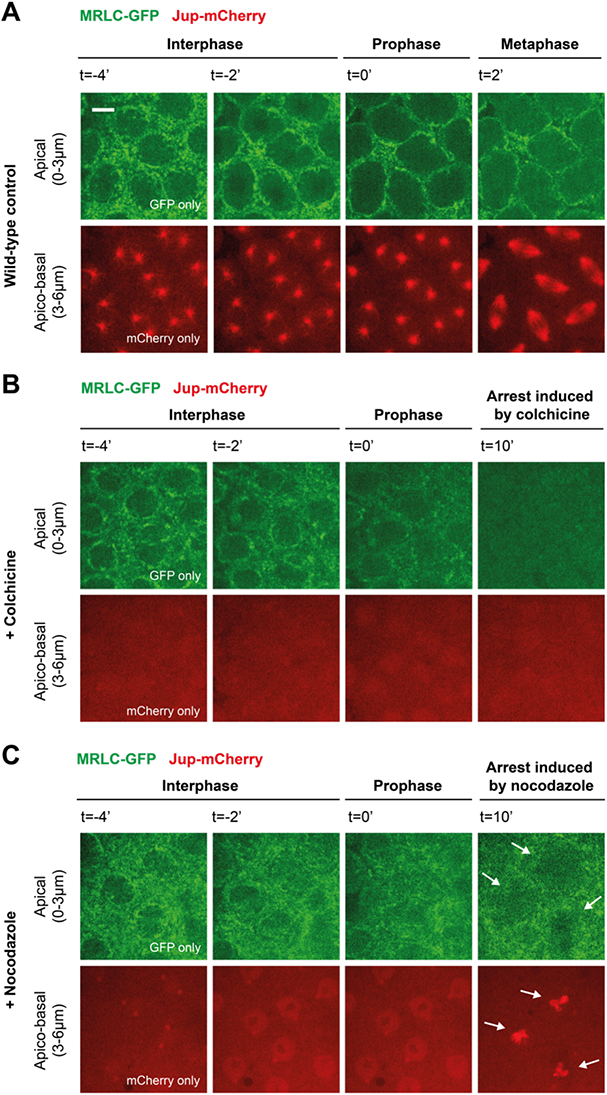
A focal pool of MTs, but not the centrosomal MTs *per se*, is required for the formation of cytoplasmic compartments. Time-lapse images depict cytoplasmic compartments (MRLC-GFP) and microtubules (Jup-mCherry) in embryos under **(A)** control conditions (n=4 embryos), or when injected with inhibitors of MT polymerization, **(B)** colchicine and **(C)** nocodazole. While colchicine injection leads to gradual destruction of cytoplasmic compartments (n=5 embryos), nocodazole injection (at ∼15ng/ml effective concentration, impairing centrosomal but not chromatin mediated MT polymerization) enables re-organization of cytoplasmic compartments, albeit less robustly (n=4 embryos). See how the beginning of chromatin-mediated MT assembly coincides with when the cytoplasmic compartments reorganize (indicated by *white* arrows). Scale bar=5μm.

**Figure S8.**
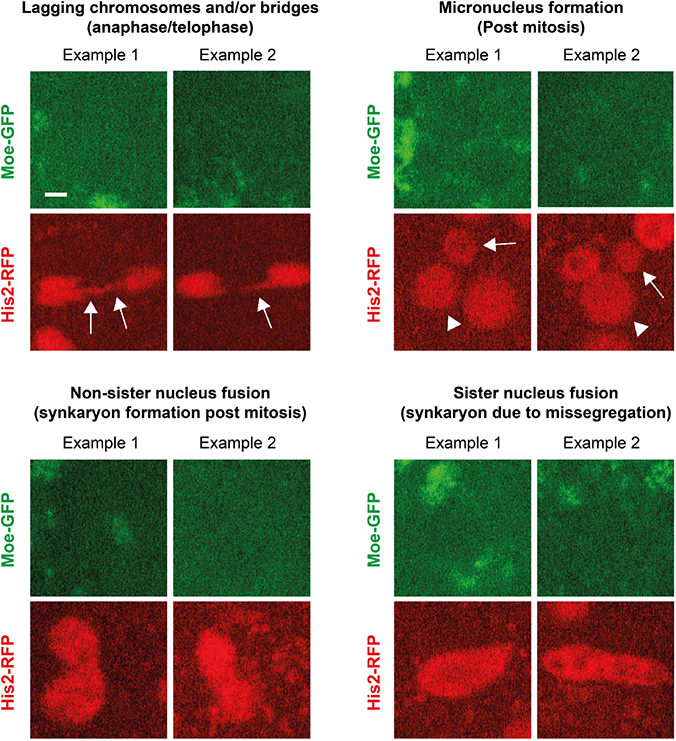
Various nuclear damages arise upon a collapse of cytoplasmic compartments. In pairs of example nuclei, micrographs illustrate four different types of nuclear damages that arise upon the collapse of cytoplasmic compartments (judged by the loss of compartment organization in the Moe-GFP channel). *White* arrows in the top left panel point out the lagging chromosomes, whereas the ones in the top right panel indicate the micronuclei that stem from their “mother” nuclei (marked by *white* arrowheads). Scale bar=2μm. Also see Supplementary text S2.

**Figure S9.**
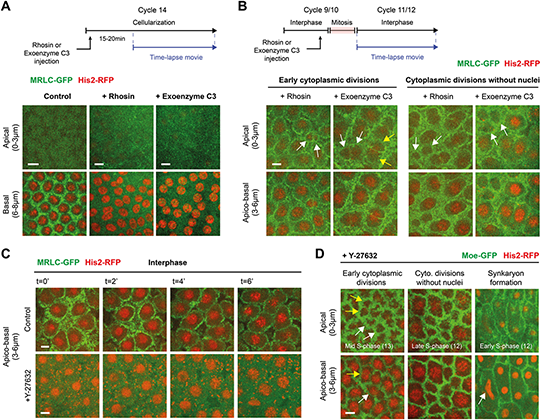
Although actin’s myosin-II based contractility is required for proper cellularization in cycle 14, it is largely dispensable for the formation and divisions of cytoplasmic compartments in the early blastoderm. **(A and B)** Micrographs illustrate myosin localization (MRLC-GFP) in control embryos or in embryos injected with RhoA-GTPase inhibitors, Rhosin or Exoenzyme C3. **(A)** In cycle 14, the process of cellularization is perturbed by the inhibition of RhoA (n=10 embryos in each condition), judged by the loss of myosin front (basal panels) and the absence of prospective cell compartments (see the lack of furrow shadows in the apical panels). **(B)** In prior cycles (10–13), however, the cytoplasmic compartments can form and divide even when RhoA is inhibited (n=5 embryos in each condition). Moreover, cytoplasmic compartments that divide in interphase (left panels; indicated by *white* or *yellow* arrows), as well as those that can divide without nuclei (right panels; indicated by *white* arrows), continued to occur in embryos where RhoA was inhibited. The schemes above panels (A and B) illustrate the injection protocols. Also see Supplementary text S3. **(C)** Time-lapse images illustrate myosin localization in embryos expressing MRLC-GFP and His2-RFP, either in control conditions (top panels) or when injected with the Rho kinase inhibitor Y-27632 (n=8 embryos). In line with previous work (*30, 60*), myosin delocalizes and redistributes to the cytoplasm upon Y-27632 injections. **(D)** In embryos injected with Y-27632 (n=8 embryos), cytoplasmic compartments can still divide in interphase (left panels; indicated by a pair of *white* or *yellow* arrows in the apical channel, with matching singlet arrows pointing at the associated nuclei in the basal channel). Similarly, the cytoplasmic divisions can continue without associated nuclei (mid-panels). Slower formation of cytoplasmic compartments in Y-276320-injected embryos (as shown in Fig. 3, F and G) occasionally leads to synkaryons (nuclear fusions) in early interphase (right panels; denoted by a *white* arrow). Scale bars=5μm. Also see Supplementary text S4.

**Figure S10.**
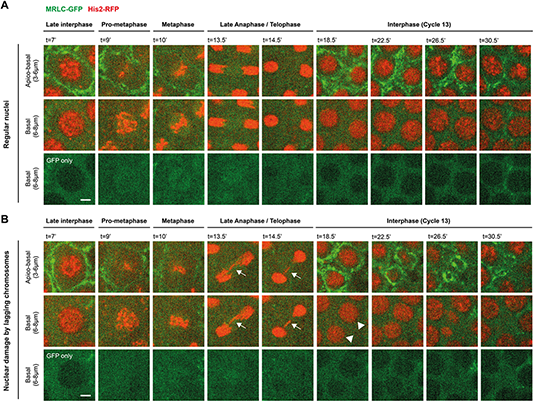
Nuclei with karyotype damages go through “nuclear fallout” in early embryos. Time-lapse micrographs show the fate of nuclei from interphase (cyc.12) to interphase (cyc. 13) in embryos expressing MRLC-GFP and His2-RFP. Basal GFP-only channels help visualizing the nuclear entry into, and exit from, mitosis. **(A)** Under normal circumstances, nuclei go through mitosis and continue to reside apically in the blastoderm. **(B)** As demonstrated by a large body of previous studies (*37–39*), nuclei with karyotype damages (e.g., lagging chromosomes that form anaphase bridges as indicated by *white* arrows) are usually eliminated from the blastoderm by fallout (follow *white* arrow heads). In our experiments, we can observe both nuclear extrusions (Fig. 4, A and B; Type 1) and fallouts (depicted here; Type 2) within the same embryos (see Video 4). Scale bars=2μm. Also see Supplementary text S5.

**Figure S11.**
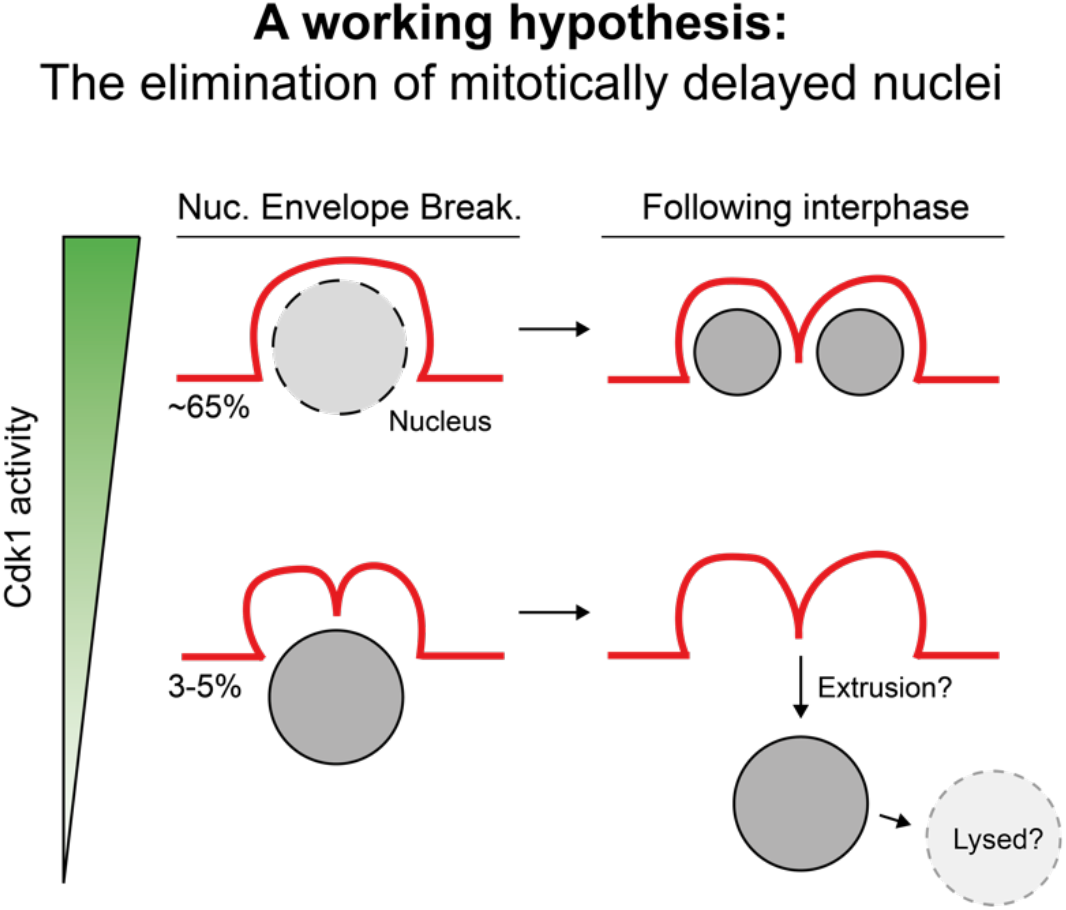
A working hypothesis on mitotically delayed nuclei and their extrusions. Cartoons illustrate our working hypothesis on the elimination of mitotically delayed nuclei in early fly embryos. As these nuclei break their envelope (NEB) late and fail to divide altogether, we hypothesized that the delayed entry is, at least in part, mediated by having less than ideal levels of Cdk1 activity (see left-hand side). As such, cytoplasmic divisions – which we found to be normally slowed by high Cdk1 activity – occur early and furrow even deeper. In the next cycle, autonomously divided cytoplasmic compartments extrude the delayed nuclei associated with them (see right-hand side). We predict that the extruded nuclei may even be lysed in the interiors of embryos. Regular NEB (top left) is indicated by a *dashed* line on the periphery of nucleus. Percentages indicate the proportion of nuclei that display the indicated phenotypes under wild-type conditions.

**Figure S12.**
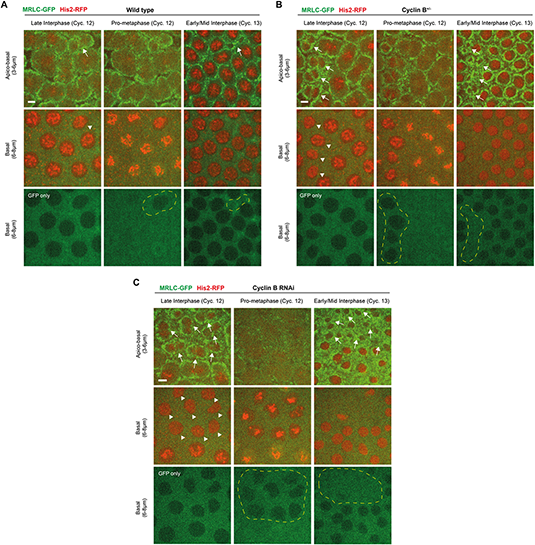
Lowering Cdk1 activity by decreasing Cyclin B levels sensitizes the cytoplasm to have more mitotically delayed nuclei and consequential extrusions. Micrographs illustrate the state of mitotically delayed nuclei (interphase to next interphase) in **(A)** control and **(B)** CycB^+/-^ embryos, or in embryos **(C)** injected with dsRNA against Cyclin B. In all conditions, the embryos express MRLC-GFP and His2-RFP simultaneously. *White* arrows (top panels) highlight the cytoplasmic compartments that divide independently of the mitotically delayed nuclei associated with them (indicated by *white* arrowheads in the basal panel). Mitotically delayed nuclei and their extrusions from the blastoderm are highlighted by *yellow dashed* lines. See the noticeable increase in the delays and extrusions in (B) and (C) in comparison to control embryos (quantifications and the sample size for each group are reported in Figure 4). Scale bars=5μm.

### Supplementary Video Captions

**Video S1. Cytoplasmic divisions normally ensue nuclear divisions in early fly embryos.** Time-lapse movie of an embryo expressing MRLC-GFP (myosin) and His2-RFP (histones), captured using a spinning-disk confocal scope through cycle 12 to the start of cycle 13. Time is reported min:sec.

**Video S2. Cytoplasmic divisions can continue for several rounds without associated nuclei in fly embryos.** Time-lapse movie of an embryo expressing MRLC-GFP (myosin) and His2-RFP (histones), captured using a spinning-disk confocal scope through mid-cycle 11 to mid-cycle 13. Example snapshots from the video are depicted in Figure 2A. Time is reported hour:min:sec.

**Video S3. Cytoplasmic compartments can form and divide without nuclei, and do this independently of the Cdk-cyclin oscillator, in arrested embryos.** Time-lapse movie of an embryo expressing MRLC-GFP and Jup-mCherry (microtubules), arrested in ∼cycle 2-3 by the injection of dsRNAs against all mitotic cyclins A-B-B3, roughly 90min prior to the capture of this video. This protocol has been used to successfully arrest embryos early in development, so as to prevent nuclear cycles at the cortex of embryos (*10, 11, 26*). As performed previously using Plk4-mNeonGreen’s retention from nuclei (*11*), the lack of nuclei was controlled by examining the MRLC-GFP channel in alternating Z-slices, as nuclei normally appear as dark circle shadows in MRCL-GFP channel in cycling embryos due to MRLC’s retention from nucleoplasm (see Fig. S4 for validations). Time is reported min:sec.

**Video S4. Nuclear cycles can continue in the absence of cytoplasmic compartments, albeit with several damages.** Time-lapse movie of an embryo expressing Moe-GFP (actin binding protein) and His2-RFP (from cycle 10 to 14), injected with cytochalasin B roughly prior to cycle 10. Although nuclear divisions can continue without cytoplasmic compartments (see their absence in Moe-GFP channel), they do so with several damages (detailed in Fig. S8). Moe-GFP channel also illustrates the intact sweeps of nuclear envelope breakdown during mitotic entry, just like in unperturbed control embryos. Also see Supplementary text S2. Time is reported min:sec.

**Video S5. Mitotic-delay associated nuclear extrusions and damage-related nuclear fallouts can occur within the same embryo.** Time-lapse movie of an embryo expressing MRLC-GFP and His2-RFP, captured through the beginning of cycle 12 to the middle of cycle 13. Mitotically delayed nuclei and their extrusions are indicated by *white* arrows labelled “Type 1”. An example nucleus with a karyotype damage (i.e., with lagging chromosomes forming an anaphase bridge) and its gradual fallout are indicated by *yellow* arrows labelled “Type 2”. MRLC-GFP channel is depicted with a single slice from basal depth, so as to help visualizing the delay of NEB for Type 1 nuclei. Also see Supplementary text S5. Time is reported min:sec.

**Video S6. Halving the genetic dose of Cyclin B sensitizes the cytoplasm for more local mitotic delays and associated nuclear extrusions.** Time-lapse movie of an embryo expressing MRLC-GFP and His2-RFP, captured through the beginning of cycle 11 to the middle of cycle 13. For ease of visualizing both mitotic delays and extrusions simultaneously, only His2-RFP channel was shown here (see Fig. S12B for both channels). Note major patches of local mitotic delays and their extrusions at large (encapsulated by *yellow* lines). The specific embryo depicted here was selected from the red data points quantified in Fig. 4F, belonging the group of embryos where we observed delays for at least >5% of nuclei in our field of view. See *Results* for our discussion on the partial genetic penetrance of CycB^+/-^ for this phenotype. Time is reported min:sec.

**Video S7. Mitotically delayed nuclei seize to be extruded in embryos where cytoplasmic compartments were abolished.** Time-lapse movie of an embryo expressing Moe-GFP (actin binding protein) and His2-RFP (from cycle 11 to 13), injected with cytochalasin B roughly prior to cycle 10. For ease of visualizing nuclear defects and eliminations, only His2-RFP channel was shown here (see Video S4 and Fig. 3E for the lack of cytoplasmic compartments under the same conditions). Note how mitotically delayed nuclei (Type 1, *white* arrows) seize to be eliminated. Meanwhile, nuclei with karyotype damages (Type 2, *yellow* arrow) continue to be eliminated (here the damage stems from sister-nucleus fusion due to incomplete abscission; also depicted in Fig. S8). See *Results* for sample size and quantifications. Also see Supplementary text S5. Time is reported min:sec.

